# Evaluation of *Mycobacterium Tuberculosis* enrichment in metagenomic samples using ONT adaptive sequencing and amplicon sequencing for identification and variant calling

**DOI:** 10.1101/2022.12.17.520855

**Authors:** Junhao Su, Wui Wang Lui, YanLam Lee, Zhenxian Zheng, Gilman Kit-Hang Siu, Timothy Ting-Leung Ng, Tong Zhang, Tommy Tsan-Yuk Lam, Hiu-Yin Lao, Wing-Cheong Yam, Kingsley King-Gee Tam, Kenneth Siu-Sing Leung, Tak-Wah Lam, Amy Wing-Sze Leung, Ruibang Luo

## Abstract

Sensitive detection of *Mycobacterium Tuberculosis* (TB) in small percentages in metagenomic samples is essential for microbial classification and drug resistance prediction, and assisting in diagnosis and treatment planning. However, traditional methods, such as bacterial culture and microscopy, are time-consuming and sometimes have limited TB detection sensitivity. Oxford Nanopore Technologies’ MinION sequencing allows rapid and simple sample preparation for whole genome and amplicon sequencing. Its recently developed adaptive sequencing selects reads from targets, while allowing real-time base-calling during sequencing to achieve sequence enrichment or depletion. Another common enrichment method is PCR amplification of the target TB genes. In this study, we compared both methods with MinION sequencing for TB detection and variant calling in metagenomic samples using both simulation runs and those with synthetic and patient samples. We found that both methods effectively enrich TB reads from a high percentage of human and other microbial DNA. We provide a simple automatic analysis framework, including quality filtering, taxonomic classification, variant calling, and antimicrobial resistance prediction, to support these detection workflows for clinical use, openly available at https://github.com/HKU-BAL/ONT-TB-NF. Depending on the patient’s medical condition and sample type (commonly including bronchoalveolar lavage fluid, blood samples, sputum, and tissues), we discuss the findings and recommend that users optimize their workflow to improve the detection limit.

## Introduction

*Mycobacterium tuberculosis* (TB) infection can be difficult to identify, and treatment delay can be deadly ^1^. The sensitivity for differentiation between the closely related *Mycobacterium tuberculosis* complex (MTBC) members varies with traditional laboratory diagnostic methods, such as staining with microscopy or PCR-based detection ^2^. Phenotypic antimicrobial susceptibility testing (AST) is commonly used for antimicrobial resistance detection, but it is time-consuming and complicated ^2,3^. In 2018, the World Health Organization (WHO) provided extensive guidelines for the use of high-throughput sequencing (including next-generation short-read sequencing and third-generation long-read sequencing) for TB detection and molecular drug susceptibility testing, with details for both targeted and metagenomic sequencing ^4^. Applying these sequencing techniques shortens the time for diagnosis, as TB cultures might not be necessary. In addition, multiplexing in most library preparation protocols can reduce the detection cost. The constraint for using metagenomic sequencing in the routine clinical diagnostic workflow is, however, a high level of human DNA contamination, as expected in clinical sputum specimens, as well as low concentration of TB in metagenomic samples ^5-7^. Also, a robust and reproducible complementary bioinformatics workflow is required for fast and accurate diagnosis.

Studies have tested the use of Oxford Nanopore Technologies (ONT) MinION sequencing for TB sequencing owing to its simple sequencing setup and the affordable long-reads generated. However, with limited throughput per MinION flowcell, it is advisable to perform TB enrichment, either by bacterial culture or PCR, to increase detection sensitivity and variant calling precision ^8^. Recently, ONT developed a selective sequencing technique to enrich or deplete target sequences controlled by software while DNA is stranded through the nanopores ^9^, where the sequencing status of each read is determined dynamically by mapping nanopore current signals ^10^ or base-called DNA bases against the target reference ^11^. In real-time sequencing, read mapping to target references can be selected (using the host DNA enrichment mode) or rejected (using the host DNA depletion mode) for further sequencing in the ONT device. The status of the DNA is determined in the first few hundred bases, and the off-target DNA strands are removed from their stranding pore. The ONT selective sequencing is initially tested for human exome sequencing and provides options for host DNA depletion in metagenomic sequencing, reducing the time required to obtain the minimum coverage per target species with some level of enrichment ^12^. Previous studies demonstrated the application of ONT adaptive sequencing for host depletion in clinical metagenomic samples for a >1.5-fold increase in coverage ^13^ and an at most five-fold increase in the yield of low-abundance species when tested with ZymoBIOMICS mock community samples ^14^. The efficiency of this software-based enrichment, however, is strongly affected by the abundance of targets, the DNA length of the library, and the computational resources available.

Compared with NGS Illumina sequencing, ONT MinION sequencing has a shorter preparation procedure and does not require large equipment maintenance, while benefiting from long reads ^15^. The capacity of multiplexing with MinION sequencing is lower and therefore requires less waiting time to acquire the minimum number of samples per batch sequence. Long-reads improve alignment accuracy and variant detection sensitivity over large repetitive regions. MinION sequencing is suitable for both native DNA and amplicon sequencing, and methylated bases can be labeled during native DNA base-calling potential to improve lineage identification and enrich AMR profiling ^16^.

In this study, we explored the efficiency of using ONT selective sequencing and PCR amplification for low-abundance TB enrichment in metagenomic samples. We tested the protocols with (1) simulation datasets, (2) synthetic metagenomic samples, and (3) clinical metagenomic samples using the portable ONT sequencing device MinION. Instead of host DNA depletion, we tested the possibility of selecting ultra-low abundance TB DNA (i.e., ∼0.1% in the metagenomic sample) from high levels of host DNA (i.e., >95% in the sample) for enrichment, i.e., host DNA enrichment, using two ONT selective sequencing toolkits (**Figure 1**). For ONT selective sequencing, we tested the performance in the whole TB genome and AMR-associated gene regions. For PCR-based enrichment testing, we followed the workflow by Tafess *et al*. ^17^, which targets 19 AMR-associated regions tested on both the Illumina and MinION platforms. We assessed the effectiveness of different strategies by the level of TB enrichment, turnaround time, and the comprehensiveness of the downstream analyses. We concluded that all the tested enrichment methods are effective in simple metagenomic samples, and that different enrichment strategies might be suitable, depending on sample properties and patient medical condition. We provide a simple, user-friendly bioinformatics workflow for TB identification after enrichment and MinION sequencing, as well as for standard drug resistance profiling.

**Figure 1.**
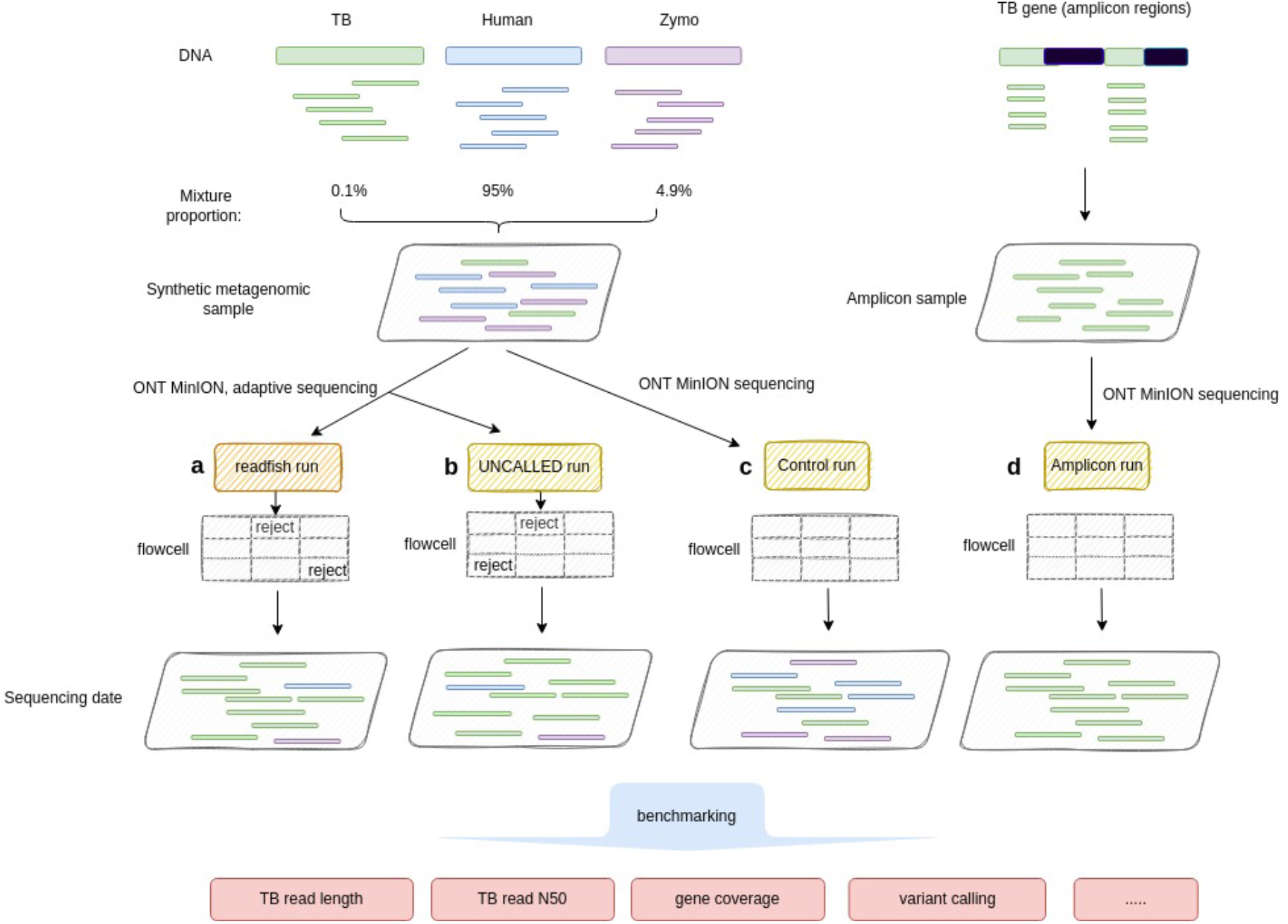
Illustration of data synthesis and data analysis. Two adaptive sequencing runs with (a) readfish, (b) UNCALLED, and (c) one control run were conducted on the synthetic metagenomic sample with 95% human DNA, 4.9% Zymo DNA, and 0.1% TB DNA. Adaptive sequencing can adaptively select or reject a sequencing read on each flowcell. (d) An Amplicon run was conducted on a selected TB gene region. All generated reads underwent benchmarking to compare different methods of enrichment of TB reads.

## Results and discussion

### Enrichment in ONT amplicon sequencing and adaptive sequencing

The number of TB reads detected in clinical samples can vary from one to thousands, with over 90% human reads ^6,7^. To simulate a simple TB metagenomic sample for enrichment evaluation using MinION amplicon sequencing and adaptive sequencing, we prepared a synthetic metagenomic sample by mixing 95% HG002 human DNA, 4.9% ZymoBIOMICS Microbial Community Standards (Zymo) DNA, and 0.1% *Mycobacterium Tuberculosis* strain H37Rv DNA (**Figure 1**). In the control sequencing run, as expected, a similar percentage of reads was recovered (92.17% human, 7.71% Zymo, and 0.12% TB reads) (**Supplementary Table 1**).

The enrichment efficiency of ONT adaptive sequencing is highly affected by the computer specifications (it is both CPU and GPU intensive), as it often requires high computational power for real-time signal processing. The reference panel size affects the speed of target selection and the accuracy of rejecting non-target reads ^11^. Also, with less repetitive and low-complexity reference sequences, the signal or read mapping quality and speed improve ^11^. To confirm the compatibility of the computing setting with the adaptive sequencing software, we recommend running some simulations of the enrichment experiment using the control dataset. We tested several software settings (i.e., no_seq, and no_map settings, **Supplementary Table 2**). We also tested the choice of reference sequences and repeated masking in the reference panel using simulation before the actual runs, and we found that there was no significant difference in using repeated masking in the reference in the enrichment.

The aim of all the tested protocols is to enrich TB DNA instead of depleting human DNA. Implementation of detection workflow is more cost-effective, especially in developing countries, if effective enrichment for identification and AMR variant calling can be achieved using one MinION flowcell. In addition to on-target coverage, the level of enrichment is associated with the sequencing yield. Since selective sequencing repetitively unblocks stranding DNA from the nanopores and might distort the structure of the pores, this reduces the total throughput of the flowcells. We found that the number of active pores decreased faster when adaptive sequencing was applied compared with no adaptive sequencing in our testing (**Supplementary Figure 1**). In addition, as a large proportion of the input DNA is non-target in adaptive sequencing, the recommended amount of DNA per MinION flowcell at traditional sequencing might be insufficient for an unknown TB percentage sample. Therefore, one possible way to improve sequencing yield during adaptive sequencing is to include more DNA. However, since limited DNA was available in most clinical samples, we restricted the use of input DNA in the benchmarking runs to the minimum required (i.e., ∼500 ng) (**Table 1**). The total throughput of the control sequencing, UNCALLED adaptive sequencing, Readfish adaptive sequencing, and amplicon sequencing using a single flowcell was 13.75 Gbp, 8.17 Gbp, 4.55 Gbp, and 12.39 Gbp, respectively (**Table 1**). After scaling with the number of available pores at the beginning of the sequencing, the adaptive sequencing runs showed an approximately 46% (readfish) and 74% (UNCALLED) reduction in total yield compared with that of the control run. Except for amplicon sequencing, the sequencing experiments were terminated only after the number of available pores dropped below 50 active pores for stranding, and therefore sequencing with base-calling alone took approximately two days. For amplicon sequencing, since the panel covers only 19 AMR-associated regions, which target mainly 267 mutations that confer resistance to 12 anti-TB drugs ^17^, it can achieve sufficient coverage per sample for variant calling within the first hour of sequencing (with average coverage of 16,255 for the target regions). Barcoding and multiplexing in a batch could improve the utilization of flowcells, but this could slow down the turnaround time because of the need to wait for sample collection.

Although the quality of flowcells vary from batch to batch, ranging from 800 to 1,600 available pores, the percentage of TB bases sequenced increased significantly by ∼3.07-fold in readfish and ∼1.98-fold in UNCALLED adaptive sequencing compared with the control run after normalization against the number of pores (**Table 1** and **Supplementary Table 1**). In the control run, only 1,974 reads (0.1% bases of total throughput) were aligned to the TB reference. The selection performance of readfish (10,834 reads; 1.15% bases of the total throughput) was better than that for UNCALLED (5,447 reads; 0.34% bases of the total throughput) with such a low abundance target (**Supplementary Table 1**). readfish enrichment allows, on average, 9.3x coverage across the TB reference genome and 9.8x coverage among the 18 AMR-associated genes (ranging from the highest coverage of 16.2x in *rpoB* to the lowest coverage of 4.7x in *rplC*) (**Supplementary Table 3**).

**Table 1.** Sequencing statistics.

| General summary | Control | readfish | UNCALLED | Amplicon | MG_TB23178 |
| --- | --- | --- | --- | --- | --- |
| Mean read length | 6,212 | 397 | 1,395 | 905 | 438 |
| Mean read quality | 13 | 13 | 13 | 13 | 10 |
| Median read length | 4,878 | 381 | 1,163 | 841 | 432 |
| Median read quality | 13 | 13 | 13 | 13 | 10 |
| Number of reads | 2,213,087 | 11,460,014 | 5,859,263 | 13,688,610 | 9,208,422 |
| Read length N50 | 10,288 | 404 | 1,842 | 976 | 464 |
| STDEV read length | 5,329 | 429 | 803 | 439 | 129 |
| Total bases | 13,746,798,479 | 4,551,882,404 | 8,170,637,866 | 12,389,957,129 | 4,033,992,524 |
| Total bases scaled by pores | 12,154,552 | 3,134,905 | 6,505,285 | 10,290,662 | 3,714,542 |
| # of primary aligned reads to TB | 1,974 | 10,834 | 5,447 | 10,602,938 | 32 |
| # of bases (Mb) aligned to TB | 10.7 | 42.0 | 23.5 | 9,172.3 | 0.03 |
| # of pores when used | 1131 | 1452 | 1256 | 1204 | 1086 |
| DNA input for sequencing (ng) | 521 | 556 | 574 | 66 | 586 |
| Sequencing duration | 2d 18h 21m | 1d 19h 13m | 1d 18h 12m | 1d 2h 46m | 2d 18h 55m |

UNCALLED enrichment allows, on average, 5.2x coverage across the TB reference genome. Compared with the average coverage of 2.4x for the whole TB genome in the control sample, readfish achieved enrichment coverage of 3.9-fold and UNCALLED achieved enrichment coverage of 2.2-fold. The higher coverage is especially important for variant calling, variant phasing, and consensus generation for AMR detection. In the amplicon sequencing, over 99.9% of the reads were assigned to TB, with only 0.03% of reads belonging to human DNA, indicating a low level of contamination (**Supplementary Table 1**). In the amplicon sequencing, there was an average of 422,642x coverage in the target regions (**Supplementary Table 4**).

Based on our testing results, selective sequencing slightly affects the read length of targeted TB. The mean and median length of the TB read decreased from 5,404.6 bp and 3,642.5 bp, respectively, in the control run to 3,876.5 bp and 1,595 bp, respectively, in the readfish run, and 4,306.8 bp and 2,474 bp in the UNCALLED run (**Supplementary Table 1**). The N50 length remained ∼9,000 bp in both the control and adaptive sequencing runs. One possible explanation might be an inaccurate assignment of TB reads as non-targets, resulting in early rejection during stranding. Therefore, we evaluated the short TB reads (i.e., below 500 bp) in adaptive sequencing runs and confirmed that the short reads did not concentrate on any particular genomic regions, suggesting that this might be due to random fragmentation during sequencing. In amplicon sequencing, the read length was constrained by the designed amplification region in the panel. The mean and N50 read length were 865 bp and 952 bp, respectively, with most of the reads sequenced as a complete amplicon without much fragmentation (**Supplementary Table 1**). Although amplicons were very specific in target enrichment, and less than 0.03% of amplicon reads were not primarily mapped to TB due to the shorter read length, 22.54% of the sequenced amplicon reads were not mapped uniquely to TB (**Table 1**).

### Removal of non-target DNA in adaptive sequencing

Since the first few hundred bases of the DNA were used to determine its identity and to decide whether the fragment should be carried on for sequencing or rejected from the standing pore in adaptive sequencing, a large proportion of the sequencing yield, which were the short fragments in our samples, were non-targets (i.e., human and Zymo reads). There were 99.76% (readfish) to 40.57% (UNCALLED) of short reads below 1000 bp in the adaptive sequencing runs, but only 14.74% of reads below 1000 bp in the control run (**Figure 2**). On the other hand, our results showed that readfish can filter noise read faster than UNCALLED. readfish’s 99% noise read had a length of less than 500 bp, while in UNCALLED, 99% of the noise reads had a length of less than 2900 bp (**Figure 2**).

**Figure 2.**
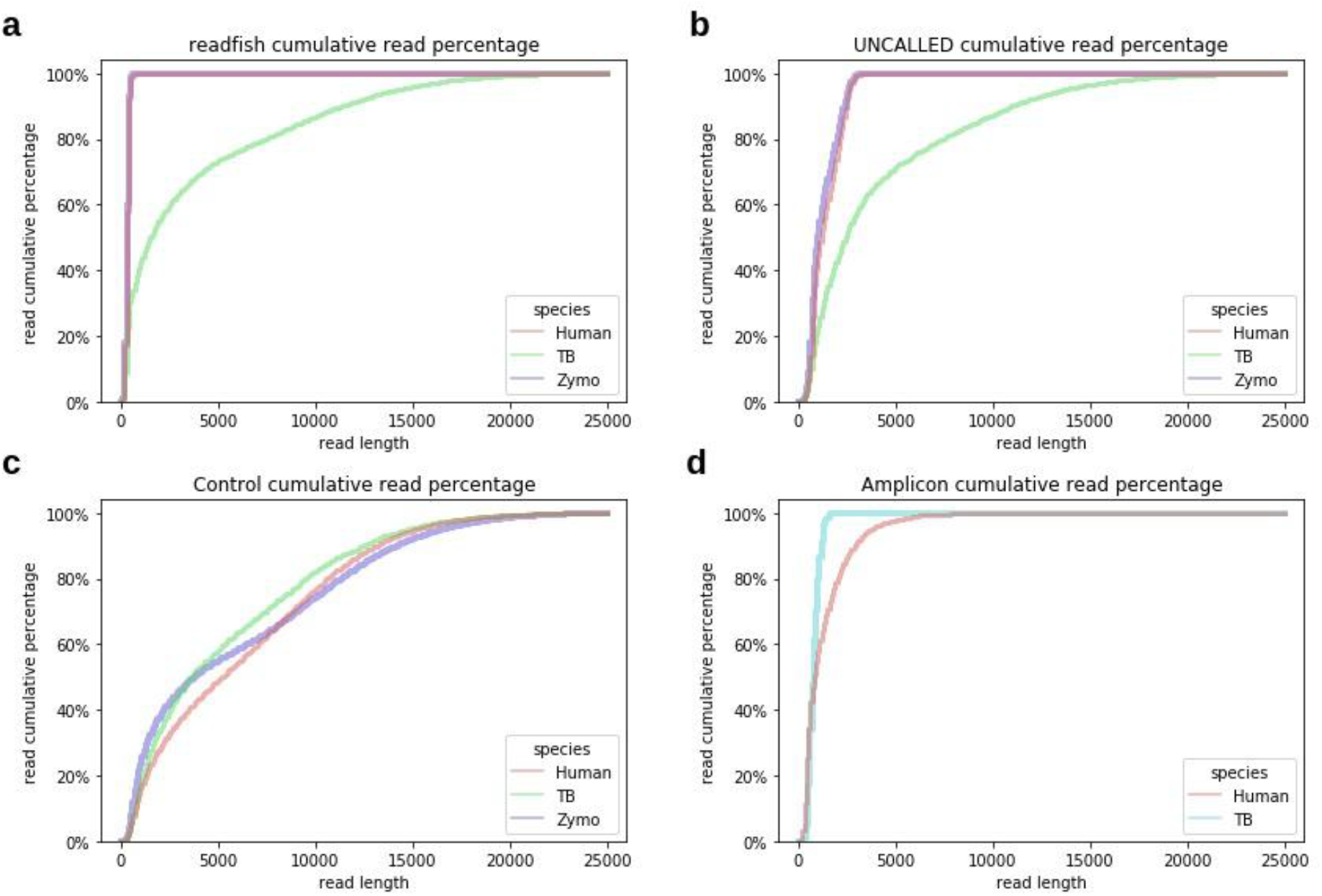
Read length distributions among different samples. The read cumulative percentage over different read lengths (bp) from different species at samples generated via adaptive sequencing of (a) a readfish run, (b) UNCALLED run, (c) Control run, and (d) Amplicon run. Compared with the control run, in which different species reads had similar read length distribution, the readfish and UNCALLED runs had a distinct pattern; the target species, TB, had a much longer read length, while the non-target reads, human and Zymo, had a limited read length (99% of human and Zymo reads <500 bp in readfish and <2900 bp in UNCALLED).

Most of these non-target short reads were human reads, while the non-target reads from the ZymoBIOMICS HMW DNA Standard species were similar in abundance, except for *Saccharomyces cerevisiae* for both readfish and UNCALLED runs. The abundance of ZymoBIOMICS HMW DNA Standard species from adaptive sequencing matched the theoretical percentage composition by genome copy of the standard sample (**Supplementary Table 1**). Previous studies using ONT adaptive sequencing on the Zymo community sample revealed that removing non-targets improves with higher molecular weight libraries, especially for high-abundance species in the sample^14^. The read length distribution was relatively optimal in our synthetic metagenomic sample for adaptive sequencing (i.e., 6,230 bp mean and 10,166 bp N50) (**Supplementary Table 1**). However, this could be a limitation for certain clinical specimens, such as sputum, which have been observed to often result in short DNA fragments. Our experimental results also suggest that the average read length and N50 of the non-target reads are significantly shorter (one-tailed t-test p-value: 3.2e-9 for read length, 1.2e-8 for N50) when using readfish compared with UNCALLED, which also suggests more effective identification and removal of non-targets during sequencing (**Figure 2**).

### Variant calling with amplicon sequencing and ONT adaptive sequencing data

Various antimicrobial resistance mechanisms, including alteration of the expression level of drug targets, production of drug inactivation enzymes, and modification of cellular structures for drug efflux, can be detected from TB gene mutations ^18^. According to the guidelines released by the WHO regarding the treatment of resistance TB and other related studies ^19,20^, SNPs, MNPs, indel mutations, and SV in at least 33 genes are known to be associated with resistance to 30 commonly applied TB treatment drugs, 17 of which were included in the tested amplicon sequencing panel.

We aligned the strand of H37Rv TB reads sequenced from the synthetic metagenomic samples to a new TB genome, NC_016804.1, to assess variant-calling performance, especially among the drug-resistance genes. The quality of variant calling was affected by the effective coverage, the per-base accuracy of the reads, and the alignment accuracy, which, in turn, were affected by the enrichment methods, the bias introduced by different sequencing platforms, and metagenomic complexity ^21^. The variant calling results for the adaptive sequencing are shown in **Figure 3**. Both the readfish and UNCALLED data shown improved the F1-score in genome-wide SNP and INDEL calling compared with the control dataset. The precision and sensitivity in SNP calling with readfish data achieved the best results at 82.85% and 91.07%, respectively. INDEL calling, however, remained suboptimal, possibly due to insufficient coverage in addition to the sequencing bias of ONT. The higher accuracy results matched the expectation described in Clair3’s paper, where variant calling accuracy is highly affected by the data coverage ^22^. The variant-calling results for the amplicon sample are shown in **Table 2**, where we compare the variant-calling results in the amplified regions. Among the 19 drug resistance-associated gene regions, all and only true variants were detected in both amplicon sequencing and adaptive-sequencing data, while the control data set shows a low level of false positives (FP) and false negatives (FN). The results suggest that both enrichment methods could improve variant calling in TB.

**Figure 3.**
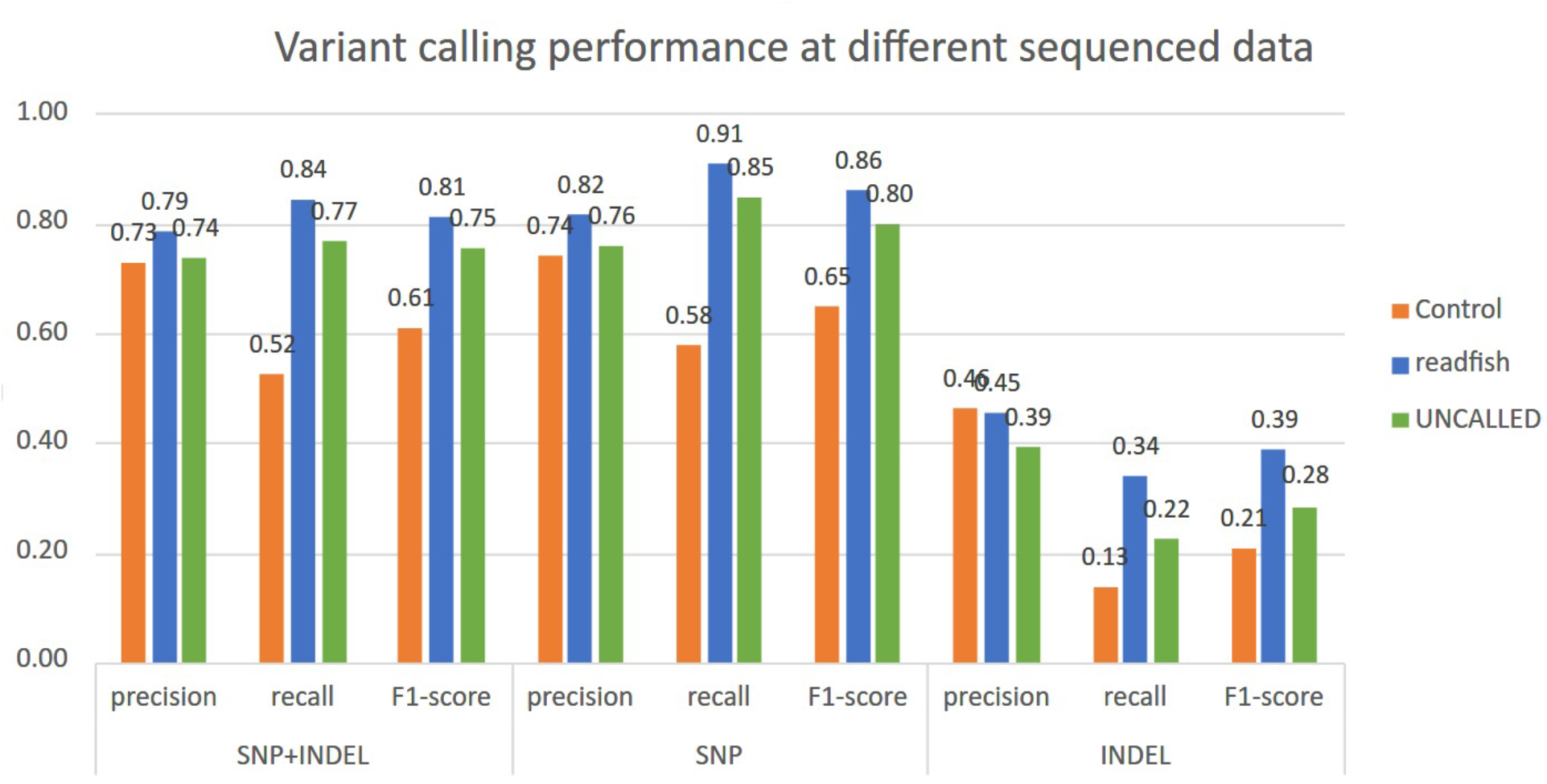
Performance of variant calling at different samples. Precision, recall and F1-score for SNP, INDEL and SNP+INDEL variants called via the Clair3v0.1-r12 guppy5 model at samples of the Control run, and adaptive sequencing run of readfish and UNCALLED.

**Table 2.** Variant-calling results in TB gene regions. The variant-calling performance of different samples at the 19 gene or locus regions defined by Tafess et al. Variant calling was conducted using the Clair3v0.1-r12 guppy5 model. TP: True Positive, FN: False negative, FP: False Positive.

| Sample | TP | FN | FP |
| --- | --- | --- | --- |
| Control | 11 | 4 | 3 |
| readfish | 15 | 0 | 0 |
| UNCALLED | 15 | 0 | 0 |
| Amplicon | 15 | 0 | 0 |

### Application of adaptive sequencing in real TB metagenomic samples

Whether using adaptive sequencing is advantageous for detecting the presence of TB or for subclonal determination depends a lot on the sample collection method, since the efficiency of adaptive sequencing is heavily affected by DNA fragment size and DNA quality. We tested the readfish protocol on a sputum sample known to have fragmented DNA based on a control Ligation sequencing kit (MG_TB23178 library) run (**Table 1**). Our results suggest a low level of enrichment, but it was not as promising as that in the synthetic sample. As a low level of enrichment leaves too many uncovered genomic positions, the sensitivity of variant calling drops drastically. This might result in a high rate of false negatives in AMR detection (i.e., 17 AMR genes do not have sufficient coverage for variant calling after readfish). In this case, amplicon sequencing, which is less sensitive to DNA fragmentation, should be considered to ensure the detection sensitivity of the test and to reduce the sequencing cost per sample.

### ONT Amplicon sequencing: Pros and cons

Amplicon sequencing works best when DNA template concentration is low and fragmented, and all variants covered within the amplicon regions are called at high confidence. This method is also commonly applied to diagnose various infectious diseases ^23^. In addition, since the throughput of a single MinION flowcell generates enough coverage data for up to 12 samples ^17^, PCR products of multiple samples can be barcoded and sequenced in batches to reduce cost and processing time.

Since the target panel size is relatively small, and the performance is predictable, there is a lower chance that the test has to be repeated owing to sequencing failure when working with low-quality samples. However, limited information is obtained from the amplification region, and it often involves tedious work for primer design and testing the amplification efficiency to change the amplification panel. Shorter amplicons generated with less specific primers from closely related taxa or contaminants can introduce ambiguity in the alignment and variant calling. While not all parts of the genome were covered, as in WGS, in this case, over 99.6% of genomic positions were not covered by the tested amplicon panel. Taxonomic classification with the amplicon data can be challenging even with longer amplicons ^24^, which is especially useful for lineage tracing ^25^. In addition, amplification efficiency among different amplicon regions might vary significantly, depending on the sequence complexity length of the amplicon and GC content ^26^. In our study, among the 19 targeted regions, *Rv0678* showed the highest amplification efficiency (with 981,074 coverage), while *rspA* showed the lowest (with 166.522 coverage) (**Supplementary Table 1**).

### A user-friendly and comprehensive bioinformatics workflow to improve turnaround time

We developed ONT-TB-NF (**Figure 4**), an easy-to-use Nextflow ^27^ pipeline to assist in processing both adaptive sequencing and amplicon ONT data for TB antibiotic-resistance detection. The pipeline has four major steps: 1) quality control with FastQC ^28^, read quality filters, and trimming with Nanofilt ^29^; 2) alignment with Minimap2 ^30^ (with map-ont mode) against the H37Rv genome; 3) genome-wide or targeted variant calling with Clair3 ^22^ with the haploid calling mode; and 4) TB-specific antimicrobial resistance prediction using TBProfiler ^31^ against the TB Profiler database. Users can start the analysis workflow directly from FAST5 files, which includes high-accuracy base-calling with Guppy 5 or FASTQ files to reduce the processing time. For amplicon date, users need to provide an additional BED file to analyze the amplicon sequencing data to specify the target regions.

**Figure 4.**
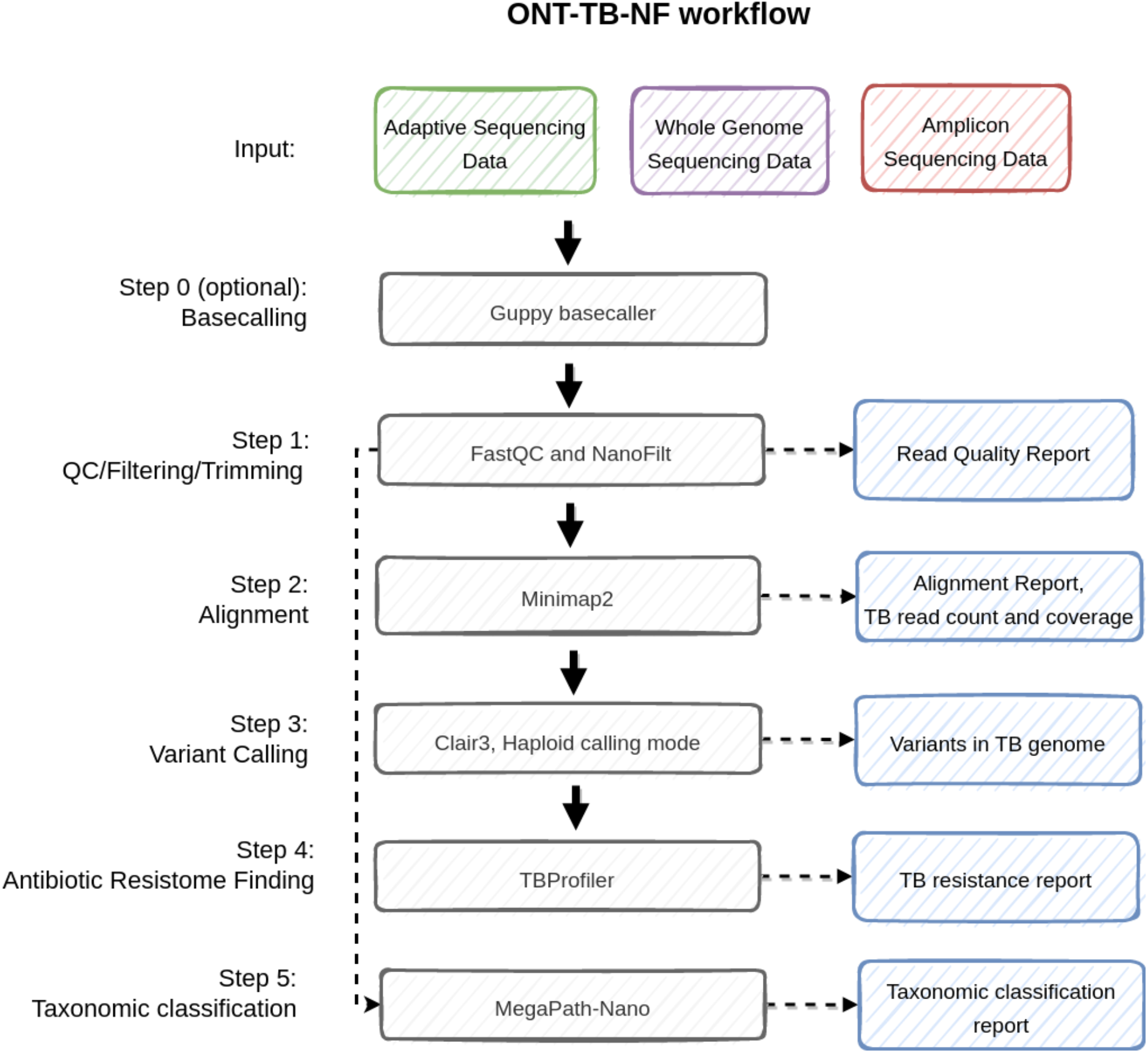
Workflow of the TB analysis pipeline.

The pipeline requires a minimal computational resource of fewer than 7GB of RAM and 36 threads. It takes approximately three minutes to process a gigabase of bases if base-calling is not needed. The turnaround time of the workflow is about 30 minutes per flowcell (estimated with 10G bases), making the workflow highly efficient for analyzing TB data. The workflow is publicly available at https://github.com/HKU-BAL/ONT-TB-NF.

## Conclusions

Target enrichment assists with accurately detecting low abundance *Mycobacterium tuberculosis* (TB) in metagenomic samples, allowing enough coverage for variant calling and antimicrobial resistance profiling. Although the benchmarking experiments are not sufficiently replicated and are limited mostly to synthetic metagenomic samples, we aimed to show a comparison between different TB enrichment methods using MinION sequencing. In this study, we demonstrated that both ONT adaptive sequencing and amplicon sequencing could effectively enrich the low abundance TB DNA in metagenomic samples. We recommend using only one sample with one MinION flowcell, as our experimental results have shown that one MinION flowcell can enrich the TB genome with an ∼9x coverage, which provides sufficient coverage to perform variant calling. While amplicon sequencing is more suitable for low-quality fragmented DNA, selective sequencing allows even whole genome enrichment and higher resolution of taxonomic classification. A different selection of enrichment methods should be considered based on the quality of specimens and the level of enrichment required. In addition, we do not expect the use of adaptive sequencing to be an effective rule-out test, especially when the bacterial load in the sample is extremely low. This is because the sensitivity of the test is significantly compromised when the sequencing coverage is too low (i.e. the enrichment level of using adaptive sequencing is not as good as using amplicon sequencing). We also provide a user-friendly workflow, ONT-TB-NF, for ONT adaptive sequencing, WGS, and amplicon data processing, to facilitate TB-specific antimicrobial resistance detection with limited computational requirements.

## Methods

In this study, we performed a total of five ONT sequencing runs: 1) a control sequencing run with a synthetic metagenomic sample, 2) a readfish adaptive sequencing run with a synthetic metagenomic sample, 3) an UNCALLED adaptive sequencing run with a synthetic metagenomic sample, 4) an amplicon sequencing run with a synthetic metagenomic sample, and 5) a readfish adaptive sequencing run with a clinical specimen (IS6110, CP=7.0).

### Preparation of the synthetic TB metagenomic sample

The synthetic metagenomic sample comprised 95% pure HG002 (Coriell Cell repositories, USA), 4.9% ZymoBIOMICS HMW DNA Standard (Zymo Research, USA), and 0.1% high molecular weight TB (NC_000962.3) DNA. According to the manufacturer’s specifications, the ZymoBIOMICS HMW DNA Standard includes *Pseudomonas Aeruginosa* (14%), *Escherichia Coli* (14%), *Salmonella Enterica* (14%), *Enterococcus Faecalis* (14%), *Staphylococcus Aureus* (14%), *Listeria Monocytogenes* (14%), *Bacillus Subtilis* (14%), and *Saccharomyces Cerevisiae* (2%). The Qubit 4.0 fluorometer (Life Technologies, USA) was used to quantify individual DNA samples before the preparation of the master mix of synthetic metagenomic samples. The synthetic metagenomic samples were stored at -20 °C after library preparation.

### Library preparation of the control and adaptive ONT sequencing runs

For each sequencing library, an input of 1 ug of DNA was quantified and fragmented to approximately 17 kb using g-TUBE (Covaris, USA) at 3500 rpm on Centrifuge 5425 (Eppendorf, Germany). The DNA was then purified and size-selected using 0.6X AMPure XP beads (Beckman Coulter, USA). The subsequent library preparation steps were performed following the ONT SQK-LSK110 Genomic DNA by ligation protocol (GDE_9108_v110_revL_10Nov2020) with the following modifications. To improve the DNA yield after end-prep, the incubation time was increased from five minutes each at 20 °C and 65 °C to 10 minutes each during the DNA repair and end-prep steps. To maximize the amount of HMW DNA recovered after each washing step, the incubation time with AMPure XP beads in all the cleaning steps and elution steps was increased from, about two to 10 minutes, to 20 minutes. After adapter ligation, approximately 50 fmol of the library was loaded into the R9.4.1 flowcell (ONT, GB) and sequenced using MinION until there were less than 50 active pores in the flowcell. The duration of each library preparation was 3 to 4.5 hours, and of each sequencing run was up to 96 hours.

### Library preparation of the amplicon sample

The library preparation was performed following the ONT SQK-LSK110 Amplicons by Ligation protocol (ACDE_9110_v110_revM_10Nov2020) using 1 ug of approximately 1000 bp amplicons. The incubation conditions applied were the same as that used for the genomic DNA. The amplicons were purified using 1X AMPure XP beads in all the cleaning steps instead. At the end of the library preparation, approximately 100 fmol of amplicon DNA was loaded and sequenced by MinION until sufficient estimated coverage was obtained. The target regions were *gyrB, gyrA, rpoB, Rv0678, rpsL, rplC, atpE, rrs, rrl, mabA-inhA, rpsA, tlyA, katG, FurA-KatG, pncA, eis, whiB7, embB*, and *ubiA*.

### Nanopore sequencing with adaptive sampling

Adaptive sequencing with readfish (0.0.6dev2) and UNCALLED (v2.2) on MinKNOW software (distribution version of 21.06.13) was used with synthetic metagenomic samples and base-called with the Guppy (v5.0.16) GPU version on a computer with two 8-core Intel i9-11900F processors and a NVIDIA GeForce RTX 2080 Ti GPU. UNCALLED was configured in “realtime” enrich mode. The TB strain of H37Rv, NC_000962.3, was used as the reference for target selection for UNCALLED. readfish was run in the targeted sequencing mode, with the mapping condition of “multi_on” and “single_on” set to “stop receiving”, and “multi_on”, “single_off”, “no_map” and “no_seq” set to “unblock”. We combined the GRCh38 human genome and the genome of the H37Rv strain, NC_000962.3, as the reference, and set the sequencing targets as “NC_000962.3” for readfish. To evaluate the performance of the fragmented clinical samples, a sputum specimen (MG_TB23178 library) was also used for readfish enrichment analysis.

### Bioinformatic analysis on the TB-enriched ONT data

To analyze the sequenced data, we first performed a quality check of the generated datasets with NanoPack ^29^ and then mapped all reads with minimap2 ^30^ (2.15-r905) to a merged reference containing human (GRCh38), TB (NC_000962.3), and the eight species listed in l ZymoBIOMICS to check the read distribution for each composed species. The alignments were filtered to remove those with an alignment score (AS) <1.2 to avoid potential mapping errors. For testing gene coverage at the amplicon sample, we gathered the 19 gene or locus regions defined by Tafess et al.^17^. For the control and adaptive sequencing samples, we tested coverage across the whole TB genome and 18 selected whole gene regions that confer resistance to anti-TB drugs: *gyrB, gyrA, rpoB, Rv0678, rpsL, rplC, atpE, rrs, rrl, inhA, rpsA, tlyA, katG, pncA, eis, whiB7, embB*, and *ubiA*. The coverage of each TB gene was computed using Mosdepth ^32^, with the target bed file provided.

### Benchmarking of variant calling on Clair3

To test variant-calling performance using different TB enrichment data, we first mapped reads from the sequencing data to another *Mycobacterium tuberculosis* strain reference sequence, NC_016804.1. The truth variants set of our synthetic sample (from NC_000962.3) was obtained by 1) mapping the original sequence of NC_000962.3 to the sequence of NC_016804.1 with “nucmer” from MUMer3 ^33^, and 2) further variant calling with “show-snps” from MUMer3 ^33^. The truth variant set contained 2362 SNPs and 327 INDELs, so it was suitable for testing variant-calling performance.

We performed Clair3 ^22^ (v0.1-r12, Guppy5 model) with the “haploid_precise” mode on all samples for variant calling. The variant-calling performance was evaluated with hap.py ^34^, and three metrics – precision, recall, and F1-score – were generated for both SNP and Indel.

## Data availability

The original sequencing outputs, fast5 files, from MinION, including the Control run, readfish run, UNCALLED run, Amplicon run, and all analysis outputs are publicly available http://www.bio8.cs.hku.hk/ont_tb.The bioinformatics workflow, ONT-TB-NF, is open-source software (BSD 3-Clause license), hosted by GitHub at https://github.com/HKU-BAL/ONT-TB-NF.

## Acknowledgments

R. L., T. Z., and T. T. Y. L. were partially supported by Hong Kong Research Grants Council grant TRS (T21-705/20-N). R. L. was partially supported by GRF (17113721), ECS (27204518), and TRS (T12-703/19-R), and the URC fund at HKU. G. K. H. S. was supported by GRF (15102220). T. T. Y. L. was supported by AIR@InnoHK funding (D24H) administered by Innovation and Technology Commission of Hong Kong Special Administrative Region.

## Additional Information

R. L. receives research funding from ONT. The remaining authors declare no competing interests.

## Author contributions

R. L. conceived the study. A. W. S. L., J. S., Y. L., and R. L. wrote the paper. T. Z., T. T. Y. L contributed to the design of the benchmarks. J. S., A. W. S. L., W. W. L., and Z. Z. analyzed the data. A. W. S. L., Y. L., G. K. H. S., T. T. L. N., H. Y. L., W. C. Y., K. K. G. T., and K. S. S. L. designed and conducted the wet experiments. W. C. Y., K. K. G. T., and K. S. S. L. collected and provided the clinical sample. All authors evaluated the results and revised the manuscript.

## Notes

http://www.bio8.cs.hku.hk/ont_tb

https://github.com/HKU-BAL/ONT-TB-NF

