## Supplementary Materials for "Evaluation of *Mycobacterium Tuberculosis* enrichment in metagenomic samples using ONT adaptive sequencing and amplicon sequencing for identification and variant calling"

### Supplementary Figures

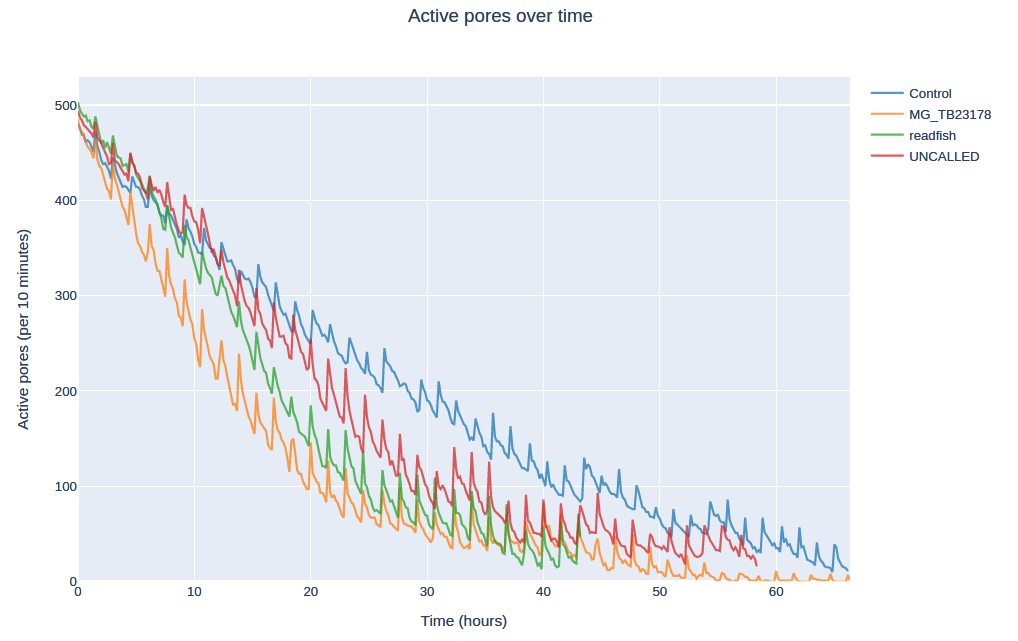

#### **Supplementary Figure 1**. Number of active pores overtime during standard sequencing or adaptive sequencing.

“MG_TB23178” is a real clinical sample. Adaptive sequencing was applied to “readfish”, “UNCALLED” and “MG_TB23178”. Standard ONT sequencing was applied to “Control”.

### Supplementary Tables

#### **Supplementary Table 1.** Read composition at different samples.

Zymo: ZymoBIOMICS Microbial Community Standards, TB: Mycobacterium tuberculosis, PA: Pseudomonas Aeruginosa, EC: Escherichia Coli, SE: Salmonella Enterica, EF: Enterococcus Faecalis, SA: Staphylococcus Aureus, LM: Listeria Monocytogenes, BS: Bacillus Subtilis, SC: Saccharomyces Cerevisiae.

| **Samples** | **Species** | **# of read** | **median** | **mean** | **N50** | **bp (Mb)** | **# of read (%)** | **bp (%)** |
| --- | --- | --- | --- | --- | --- | --- | --- | --- |
| Control | Human | 1,517,867 | 5,259 | 6,238.62 | 10,072 | 9,469.4 | 92.17 | 92.29 |
|  | TB | 1,974 | 3,643 | 5,404.63 | 9,434 | 10.7 | 0.12 | 0.10 |
|  | Zymo | 127,005 | 3,903 | 6,144.28 | 11,442 | 780.4 | 7.71 | 7.61 |
|  | All | 1,646,846 | 5,179 | 6,230.35 | 10,166 | 10,260.4 | 100.00 | 100.00 |
|  | PA | 9,882 | 8,954 | 8,889.59 | 12,163 | 87.9 | 0.60 | 0.86 |
|  | EC | 31,478 | 1,222 | 3,422.46 | 10,129 | 107.7 | 1.91 | 1.05 |
|  | SE | 31,803 | 1,879 | 3,496.56 | 6,861 | 111.2 | 1.93 | 1.08 |
|  | EF | 15,389 | 7,217 | 7,628.75 | 11,516 | 117.4 | 0.93 | 1.14 |
|  | SA | 14,151 | 9,179 | 8,952.95 | 12,393 | 126.7 | 0.86 | 1.23 |
|  | LM | 11,881 | 9,709 | 9,459.71 | 12,458 | 112.4 | 0.72 | 1.10 |
|  | BS | 10,798 | 9,713 | 9,493.50 | 12,465 | 102.5 | 0.66 | 1.00 |
|  | SC | 1,623 | 9,103 | 8,983.22 | 12,486 | 14.6 | 0.10 | 0.14 |
| readfish | Human | 8,505,147 | 383 | 393.78 | 404 | 3,349.2 | 92.35 | 91.46 |
|  | TB | 10,834 | 1,595 | 3,876.52 | 9,791 | 42.0 | 0.12 | 1.15 |
|  | Zymo | 693,409 | 378 | 390.23 | 399 | 270.6 | 7.53 | 7.39 |
|  | All | 9,209,390 | 382 | 397.61 | 405 | 3,661.8 | 100.00 | 100.00 |
|  | PA | 55,211 | 375 | 391.09 | 396 | 21.6 | 0.60 | 0.59 |
|  | EC | 166,765 | 374 | 380.16 | 393 | 63.4 | 1.81 | 1.73 |
|  | SE | 172,930 | 375 | 379.74 | 394 | 65.7 | 1.88 | 1.79 |
|  | EF | 87,058 | 384 | 401.26 | 406 | 34.9 | 0.95 | 0.95 |
|  | SA | 78,688 | 385 | 401.67 | 408 | 31.6 | 0.85 | 0.86 |
|  | LM | 64,547 | 383 | 403.50 | 407 | 26.0 | 0.70 | 0.71 |
|  | BS | 58,574 | 380 | 400.68 | 403 | 23.5 | 0.64 | 0.64 |
|  | SC | 9,636 | 384 | 402.61 | 407 | 3.9 | 0.10 | 0.11 |
| UNCALLED | Human | 4,402,901 | 1,235 | 1,435.06 | 1,893 | 6,318.4 | 92.43 | 92.73 |
|  | TB | 5,447 | 2,474 | 4,306.84 | 8,472 | 23.5 | 0.11 | 0.34 |
|  | Zymo | 355,144 | 1,084 | 1,329.37 | 1,733 | 472.1 | 7.46 | 6.93 |
|  | All | 4,763,492 | 1,223 | 1,430.46 | 1,887 | 6,814.0 | 100.00 | 100.00 |
|  | PA | 27,149 | 1,332 | 1,489.68 | 1,925 | 40.4 | 0.57 | 0.59 |
|  | EC | 88,123 | 841 | 1,087.10 | 1,269 | 95.8 | 1.85 | 1.41 |
|  | SE | 94,381 | 971 | 1,218.28 | 1,493 | 115.0 | 1.98 | 1.69 |
|  | EF | 43,141 | 1,293 | 1,484.52 | 1,925 | 64.0 | 0.91 | 0.94 |
|  | SA | 37,785 | 1,342 | 1,521.47 | 1,990 | 57.5 | 0.79 | 0.84 |
|  | LM | 31,291 | 1,377 | 1,538.22 | 1,995 | 48.1 | 0.66 | 0.71 |
|  | BS | 28,626 | 1,378 | 1,539.56 | 1,987 | 44.1 | 0.60 | 0.65 |
|  | SC | 4,648 | 1,396 | 1,539.49 | 2,003 | 7.2 | 0.10 | 0.11 |
| Amplicon | Human | 2,852 | 884 | 1,355.76 | 2,091 | 3.9 | 0.03 | 0.04 |
|  | TB | 10,602,938 | 832 | 865.07 | 952 | 9,172.3 | 99.97 | 99.95 |
|  | All | 10,605,790 | 832 | 865.20 | 952 | 9,176.1 | 100.00 | 100.00 |
| MG_TB23178 | Human | 5,037,611 | 405 | 410.56 | 436 | 2,068.22 | 100.00 | 100.00 |
|  | TB | 32 | 464 | 862.94 | 1,516 | 0.03 | 0.00 | 0.00 |
|  | All | 5,037,643 | 405 | 410.56 | 436 | 2,068.25 | 100.00 | 100.00 |

#### **Supplementary Table 2.** Simulation of different adaptive sequencing settings.

| **Setting/results** | **Setting 1** | **Setting 2** |
| --- | --- | --- |
| mode | readfish enrichment | readfish enrichment |
| host sequences | GRCh38 + NC_000962.3 | GRCh38 + NC_000962.3 |
| targets | NC_000962.3 | NC_000962.3 |
| no_seq | unblock | process |
| no_map | unblock | process |
| single_on | stop_receiving | stop_receiving |
| multip_on | stop_receiving | stop_receiving |
| single_off | unblock | unblock |
| multip_off | unblock | unblock |
| # of TB read | 18 | 16 |
| mean TB read length | 4871 | 3509 |
| TB N50 | 21135 | 20609 |
| # of human read | 128096 | 128192 |
| mean human read length | 407.3 | 407.3 |
| human N50 | 423.0 | 423.4 |

#### **Supplementary Table 3.** TB gene coverage in different samples.

| **Reference** | **Regions** | | | **Original Coverage** | | | |
| --- | --- | --- | --- | --- | --- | --- | --- |
|  |  |  |  | **Control** | **readfish** | **UNCALLED** | **Amplicon** |
| NC_000962.3 | whole genome | | | 2.36 | 9.3 | 5.2 | NA |
| NC_000962.3 | 5240 | 7267 | *gyrB* | 3.94 | 9.75 | 4.7 | 212,317.70 |
| NC_000962.3 | 7302 | 9818 | *gyrA* | 3.45 | 7.1 | 6.12 | 191,954.48 |
| NC_000962.3 | 759807 | 763325 | *rpoB* | 2.17 | 16.21 | 5.24 | 129,634.12 |
| NC_000962.3 | 778990 | 779487 | *Rv0678* | 1.55 | 11.42 | 7.91 | 893,276.92 |
| NC_000962.3 | 781560 | 781934 | *rpsL* | 3.91 | 8.83 | 5.82 | 720,359.48 |
| NC_000962.3 | 800809 | 801462 | *rplC* | 2.41 | 4.66 | 1.99 | 860,951.25 |
| NC_000962.3 | 1461045 | 1461290 | *atpE* | 2.94 | 12.28 | 1.3 | 861,528.96 |
| NC_000962.3 | 1471846 | 1473382 | *rrs* | 1.99 | 13.02 | 5.05 | 239,816.10 |
| NC_000962.3 | 1473658 | 1476795 | *rrl* | 3.05 | 12.9 | 5.95 | 134,403.83 |
| NC_000962.3 | 1674202 | 1675011 | *inhA* | 3.35 | 11.42 | 4.85 | 135,781.81 |
| NC_000962.3 | 1833542 | 1834987 | *rpsA* | 4.21 | 10.36 | 4.17 | 166,901.77 |
| NC_000962.3 | 1917940 | 1918746 | *tlyA* | 0.95 | 9.54 | 4.76 | 595,159.24 |
| NC_000962.3 | 2153889 | 2156111 | *katG* | 1.73 | 6.67 | 7.09 | 195,592.47 |
| NC_000962.3 | 2288681 | 2289241 | *pncA* | 6.66 | 7.3 | 3.64 | 664,184.18 |
| NC_000962.3 | 2714124 | 2715332 | *eis* | 3.03 | 8.88 | 6.33 | 308,059.90 |
| NC_000962.3 | 3568401 | 3568679 | *whiB7* | 1.68 | 7.3 | 0 | 727,769.09 |
| NC_000962.3 | 4246514 | 4249810 | *embB* | 2.16 | 11.92 | 3.87 | 165,897.88 |
| NC_000962.3 | 4268925 | 4269833 | *ubiA* | 2.06 | 7.38 | 6.25 | 403,959.49 |
| NC_000962.3 | AMR genes median | | | 2.06 | 7.38 | 6.25 | 403,959.49 |
| NC_000962.3 | AMR genes mean | | | 2.85 | 9.83 | 4.72 | 422,641.59 |
| NC_000962.3 | AMR genes standard deviation | | | 1.32 | 2.88 | 2.02 | 292,507.34 |

#### **Supplementary Table 4.** Coverage of the 19 amplicon sequencing regions in different samples.

| **Reference** | **Regions** | | | **Original Coverage** | | | |
| --- | --- | --- | --- | --- | --- | --- | --- |
|  |  |  |  | **Control** | **readfish** | **UNCALLED** | **Amplicon** |
| NC_000962.3 | 5958 | 6969 | gyrB | 3.93 | 9.62 | 4.78 | 425,112.07 |
| NC_000962.3 | 7388 | 8095 | gyrA | 3.95 | 9.48 | 4.67 | 682,120.97 |
| NC_000962.3 | 760076 | 761389 | rpoB | 2.51 | 15.79 | 6.33 | 347,009.81 |
| NC_000962.3 | 778924 | 779513 | Rv0678 | 1.5 | 11.42 | 7.9 | 891,074.55 |
| NC_000962.3 | 781529 | 782000 | rpsL | 3.9 | 8.74 | 5.9 | 717,429.06 |
| NC_000962.3 | 800811 | 801476 | rplC | 2.42 | 4.65 | 1.99 | 861,739.55 |
| NC_000962.3 | 1461012 | 1461437 | atpE | 2.93 | 12.42 | 1.31 | 856,858.94 |
| NC_000962.3 | 1472290 | 1473458 | rrs | 1.99 | 13.38 | 4.69 | 336,659.09 |
| NC_000962.3 | 1475575 | 1476633 | rrl | 3.64 | 13.44 | 6.88 | 398,151.60 |
| NC_000962.3 | 1673292 | 1674532 | mabA-inhA | 3.94 | 12.53 | 5.17 | 331,163.91 |
| NC_000962.3 | 1833451 | 1835007 | rpsA | 4.3 | 10.32 | 4.12 | 166,522.34 |
| NC_000962.3 | 1917862 | 1918762 | tlyA | 0.88 | 9.56 | 4.68 | 593,787.05 |
| NC_000962.3 | 2154044 | 2155443 | katG | 1.81 | 5.55 | 6.89 | 181,217.25 |
| NC_000962.3 | 2155505 | 2156396 | FurA-KatG | 1.47 | 10.07 | 7.57 | 297,863.92 |
| NC_000962.3 | 2288651 | 2289419 | pncA | 6.39 | 7.4 | 3.69 | 662,905.19 |
| NC_000962.3 | 2714908 | 2715456 | eis | 2.97 | 9.83 | 6.46 | 873,868.65 |
| NC_000962.3 | 3568312 | 3569115 | whiB7 | 2.11 | 8.16 | 0 | 723,702.49 |
| NC_000962.3 | 4247152 | 4248111 | embB | 2.07 | 11.79 | 2.98 | 569,647.89 |
| NC_000962.3 | 4268836 | 4269954 | ubiA | 2.15 | 7.43 | 6.45 | 402,008.98 |
| NC_000962.3 | AMR genes median | | | 2.06 | 7.38 | 6.25 | 403,959.49 |
| NC_000962.3 | AMR genes mean | | | 2.85 | 9.83 | 4.72 | 422,641.59 |
| NC_000962.3 | AMR genes standard deviation | | | 1.32 | 2.88 | 2.02 | 292,507.34 |
